# Live-cell imaging uncovers the relationship between histone acetylation, transcription initiation, and nucleosome mobility

**DOI:** 10.1101/2023.03.02.530854

**Authors:** Matthew N. Saxton, Tatsuya Morisaki, Diego Krapf, Hiroshi Kimura, Timothy J. Stasevich

## Abstract

Post-translational protein modifications play an important role in the regulation of gene dynamics. Certain modifications, such as histone acetylation and RNA polymerase II phosphorylation, are associated with transcriptionally active chromatin. However, the spatial and temporal relationship between chromatin and post-translational protein modifications, and how these dynamics facilitate selective gene expression, remain poorly understood. In this study, we address this problem by developing a general methodology for quantifying in live cells the dynamics of chromatin across multiple time and length scales in the context of residue-specific protein modifications. By combining Fab-based labeling of endogenous protein modifications with single-molecule imaging, we track the dynamics of chromatin enriched with histone H3 Lysine-27 acetylation (H3K27ac) and RNA polymerase II Serine-5 phosphorylation (RNAP2-Ser5ph). Our analysis reveals chromatin enriched with H3K27ac is separated from chromatin enriched with RNAP2-Ser5ph. Furthermore, in these separated sites, we show the presence of the two modifications are inversely correlated with one another on the minutes timescale. We then track single nucleosomes in both types of sites on the sub-second timescale and again find evidence for distinct and opposing changes in their diffusive behavior. While nucleosomes diffuse ∼15% faster in chromatin enriched with H3K27ac, they diffuse ∼15% slower in chromatin enriched with RNAP2-Ser5ph. Taken together, these results argue that high levels of H3K27ac and RNAP2-Ser5ph are not often present together at the same place and time, but rather each modification marks distinct sites that are transcriptionally poised or active, respectively.

## Introduction

Transcription in eukaryotic cells occurs in the context of nucleosomes, the fundamental unit of chromatin. Nucleosomes are composed of a strand of DNA wrapped around an octet of core histones^1^. This wrapped configuration provides an additional layer of gene regulation as it limits the accessibility of DNA to transcription machinery^2^. To make genes more or less accessible, the tails of histones are post-translationally modified at a large number of unique residues. These post-translational modifications (PTMs) – which include acetylation, methylation, ubiquitination, and phosphorylation, among others – change the local chemical environment of chromatin to alter the ease by which nucleosomes can slide across DNA and interact with one another^3,4^. They also create binding substrates for chromatin remodelers or transcription factors. Collectively, histone PTMs are thought to create a ‘code’ or ‘language’ that fine-tunes chromatin spatio-temporal dynamics^3–5^.

Besides histones, the transcription machinery itself is also subject to a variety of PTMs that regulate gene expression. A critical example is the phosphorylation of RNA polymerase II (RNAP2) – the complex responsible for the transcription of the vast majority of protein-coding genes. RNAP2 is phosphorylated at key residues along a conserved heptad sequence that is repeated (∼52 times in humans) within the C-terminus of the large, catalytic RPB1 subunit. Phosphorylation occurs sequentially and is thought to delineate major events in the RNAP2 transcription cycle^6–9^. In general, similar to histone PTMs, RNAP2-specific PTMs create binding substrates to recruit specific factors that facilitate transcription. As an additional regulatory mechanism, RNAP2 phosphorylation alters the local chemical environment, leading to functionally distinct clusters^10–12^.

Histone and RNAP2 PTMs are tightly coupled during gene activation. Histone acetylation at Lysine 27 (H3K27ac), in particular, is strongly correlated with RNAP2 Serine 5 phosphorylation (RNAP2-Ser5ph), a marker of transcription initiation and pausing at promoters and enhancers^13,14^. A wealth of classic biofractionation, immunoprecipitation, and structural studies have led to a standard textbook model for this correlation^3,15–17^. In the model, acetylation neutralizes the positive histone charge and reduces hydrogen bond formation. These processes weaken interactions with negatively charged DNA and other nucleosomes, so individual nucleosomes can move more easily. Acetylation also creates binding substrates for chromatin-remodelers and other trans-factors with acetyl-binding domains^18^. The end result is decondensed, loose, and more mobile chromatin where enhancers and promoters can more easily come into contact and RNAP2 can more easily be recruited and initiated for efficient transcription. Super-resolution imaging of histones in intact, fixed cells further support this model of chromatin decondensation by histone acetylation^19^, and numerous genome-wide studies have confirmed acetylation is co-enriched with RNAP2-Ser5ph near gene and enhancer transcription start sites^7,14^.

Although the textbook model provides a clear snapshot of the impact histone acetylation has on chromatin and transcription, the dynamics underlying the model remain poorly understood and have not been validated. This gap arises because PTM dynamics are hard to capture experimentally. Histone acetylation is a dynamic mark whose levels can rapidly change through the fine tuning of the activities of lysine deacetylases (KDAC) and acetyltransferases (KHAT)^15,20^. RNAP2 phosphorylation also rapidly changes as RNAP2 progresses through the various stages of the transcription cycle^7,8^. Resolving the spatiotemporal relationships between highly dynamic acetylation and phosphorylation is therefore especially challenging. Current knowledge of PTM relationships has therefore relied on fixed cell assays, including chromatin immunoprecipitation, immunostaining, and western blotting. However, in fixed-cell experiments, the dynamics are blurred by the experimental necessity of fixation (which mixes signals in time) and, in many cases, cell population averaging (which mixes signals in space).

To better resolve PTM dynamics, several live-cell imaging techniques have been developed over the last decade. First, we and others have developed technology to directly image and quantify endogenous histone and RNAP2 PTMs in living cells^21–26^. In these studies, fluorescent antibody-based probes are used to rapidly bind and light up residue-specific PTMs in distinct colors. This technology makes it possible to record the temporal fluctuations of the various modifications and better understand their dynamic relationships. Second, individual genetic loci, genes, and nucleosomes can now be tracked in living cells, making it possible to directly measure their mobilities. For example, DNA fluorescence amplification tags can be used to track single genetic loci^27–30^, RNA fluorescence amplification tags can be used to track single-gene transcription^31^, and both DNA and RNA amplification tags can be combined^32–34^. More recently, advances in single-molecule fluorescence microscopy^35,36^ and fluorescent dye development^37^ have made it possible to track and quantify the mobility of single nucleosomes^35,38–42^. Together, these new technologies are revealing a high degree of heterogeneity in chromatin mobility, both within and across cells.

Of the various live-cell imaging studies to date, discrepancies have begun to emerge that raise several fundamental questions about the dynamic relationship between chromatin, gene activation, and histone acetylation. Although there seems to be consensus that dense, constitutive heterochromatin is less mobile than average^38,39,41,43^, the mobility of less dense euchromatin remains debatable. According to the transcription ‘factory’^44,45^ or ‘hub’^46^ model, the transcription machinery is thought to be relatively immobile, causing nearby chromatin to be constrained or anchored. Recent support for this model comes from several studies that tracked specific genetic loci in living cells and showed their mobility goes down when associated with transcription^32–34,47^. Additional support comes from single-nucleosome tracking experiments in which a variety of global inhibitors and perturbations of transcription were used to show transcription generally confines single-nucleosome movements^38,48^. Nevertheless, direct imaging of RNAP2^49^ and other components of transcriptional hubs^50,51^ has revealed they are highly dynamic and transient structures, leaving it questionable to what degree they can immobilize chromatin. Furthermore, some genes and regulatory elements near genes have been observed to become *more* dynamic upon transcription, providing support for an opposing model whereby transcription ‘stirs up’ chromatin^52^, in opposition to the factory concept. Finally, the same single-nucleosome tracking studies that showed transcription slows down chromatin also showed generalized acetylation speeds it up^38^. These results lead to a paradox: if histone acetylation and transcription initiation act together at the same place and time in the nucleus, as the standard textbook model predicts, how can active chromatin be both more and less mobile?

Here we directly confront this conundrum by combining single molecule tracking^35,36^ with Fab-based imaging of live-endogenous modifications (FabLEM)^21,22,24,53^. The combination of these technologies allow us to simultaneously label and directly monitor (1) chromatin dynamics, (2) transcription initiation (RNAP2-Ser5ph), and (3) histone H3 Lysine 27 acetylation (H3K27ac), all without the use of broad-acting inhibitors or perturbations. Using this approach, we show that chromatin enriched with H3K27ac is for the most part physically separated from chromatin enriched with RNAP2-Ser5ph. We furthermore show that the two modifications localize to regions that are oppositely correlated with one another and the mobilities of nucleosomes therein are substantially different. Taken together, our data support a model wherein histone acetylation and transcription initiation are enriched in functionally separate chromatin regions with distinct physical behavior.

## Results

### Monitoring global chromatin dynamics in the context of histone acetylation and RNAP2 phosphorylation

To quantitatively explore the relationship between histone acetylation, RNAP2 phosphorylation, and chromatin dynamics, we first created an RPE1 cell line that stably expresses Halo-tagged histone H2B (H2B-Halo; **Sup. Fig. 1A**). With this cell line, we investigated the dynamics of chromatin at multiple time and length scales depending on the concentration of added Halo ligand. At higher concentrations, we could label a significant fraction of H2B, allowing us to measure the dynamics of subcellular chromatin regions on the minutes to hours timescale (**Sup. Fig. 1B**); At lower concentrations, we could label a tiny subset of H2B, allowing us to measure the dynamics of individual nucleosomes on the sub-second to minutes timescale (**Sup. Fig. 1C**,**D**).

To place these measurements within the local context of histone acetylation and RNAP2 phosphorylation, we used FabLEM^23^ to co-image endogenous histone H3 Lysine 27 acetylation (H3K27ac) – marking active genes and enhancer DNA^4,54–56^ – and endogenous RNAP2 Serine 5 phosphorylation (RNAP2-Ser5ph) – marking transcription initiation sites^8^. We were motivated by our previous experiments in which we imaged the same modifications at an artificial tandem gene array activated by the synthetic hormone dexamethasone^22^. One of our main findings was that H3K27ac can facilitate gene activation and chromatin decondensation, leading to more efficient RNAP2 recruitment and promoter escape. Due to the artificial nature of the tandem gene array, however, we worried about the generalizability of our findings and wondered if more natural and less repetitive chromatin would behave the same.

To directly test this hypothesis, we co-loaded our stable H2B-Halo cells with the same Fab (fragmented antigen binding regions) we used in the earlier study and performed 3-color confocal time lapse microscopy. As illustrated in **Fig. 1A**, we imaged endogenous H3K27ac using an AF488-conjugated Fab (green), endogenous RNAP2-Ser5ph using a CF640-conjugated Fab (purple), and H2B-Halo using JFX-554-labeled Halo ligand^57^ (orange). Unlike standard fluorescent protein fusion tags like GFP, which are permanently attached, Fabs only bind their targets transiently. This reduces interference with the underlying biology and allows Fabs to rapidly respond to changes in the post-translational protein modification landscape^24,53^. To further minimize interference, we kept the concentration of Fab relatively low in cells so target modifications were far from saturated, as we had in our earlier study^22^. By imaging cells every 2-3 minutes for 100 time points total (representing 200 or 300 minutes, respectively), we generated a complementary pair of datasets that we could mine to precisely quantify the spatiotemporal relationship between H3K27ac and RNAP2-Ser5ph across the entire nucleus (**Fig. 1B**).

**Fig. 1:**
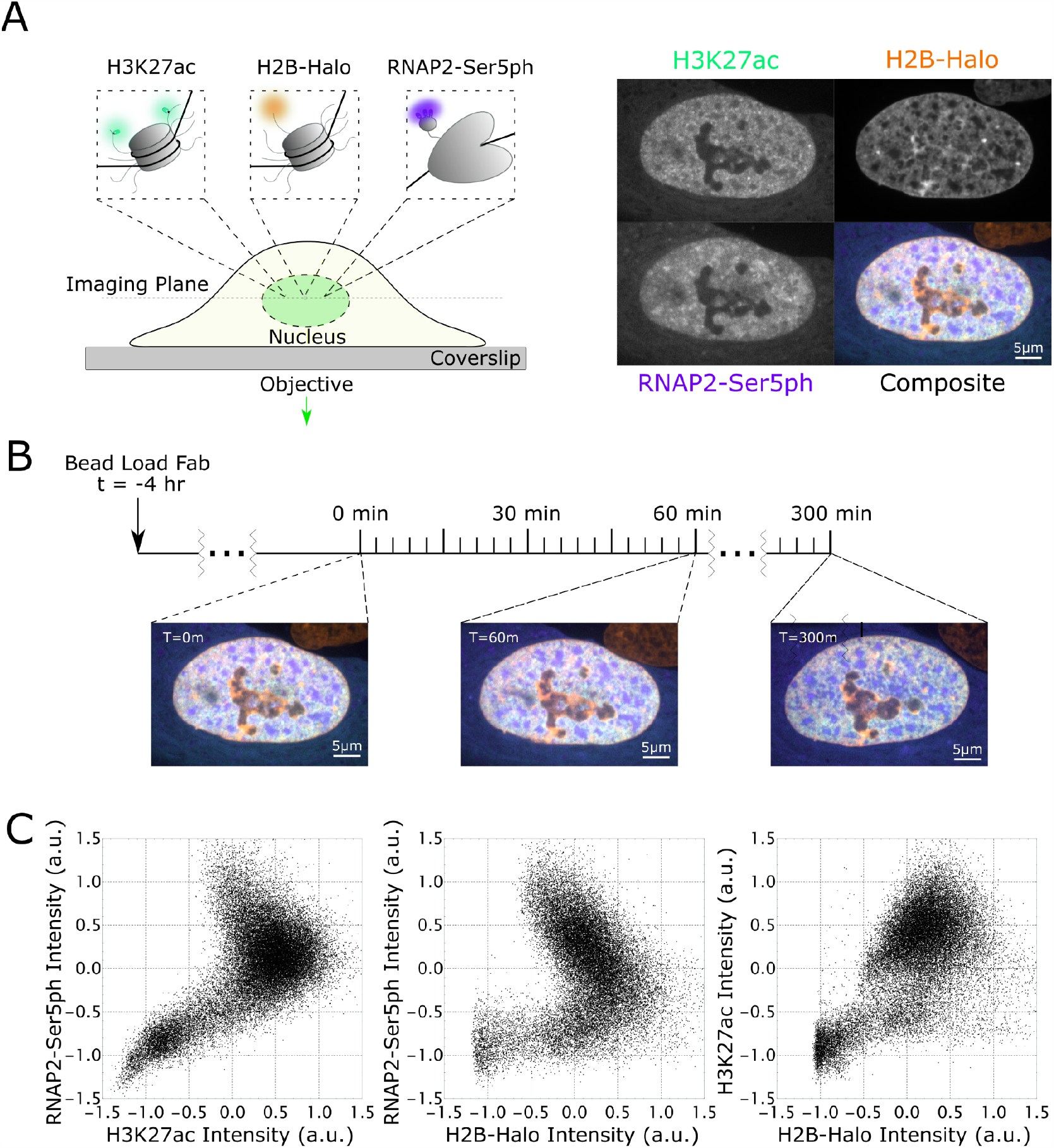
Simultaneous imaging of RNAP2-Ser5ph, H2B, and H3K27ac in living RPE1 cells. **A.** Left) Schematic of imaging system. H2B-Halo is stably expressed in RPE1 cells and stained with Halo-JFX-554 ligand. AF488-H3K27ac and CF640-RNAP2-Ser5ph specific Fab are beadloaded into cells, marking each modification in real-time. Right) Sample images and composite for all channels. **B.** Experimental time course. Cells are beadloaded with Fab 4 hours prior to imaging, and then H3K27ac, RNAP2-Ser5ph, and H2B channels are collected in a single axial z-plane every 3 minutes for 300 minutes. **C.** Scatterplots of renormalized H3K27ac, RNAP2-Ser5ph and H2B signal intensities. Each plotted point is the paired intensity values of the same pixel inside of a single nucleus at a single time point, with 3 representative plots chosen, one for each pairing of channels.

To begin to quantify these data in a simple manner, we asked to what degree the H3K27ac and RNAP2-Ser5ph signals were spatially correlated. On the one hand, standard immunoprecipitation assays have long observed a strong positive correlation between histone acetylation and transcription^15,58^. According to this body of work, genes with higher H3K27ac levels should have higher RNAP2-Ser5ph levels^4^. On the other hand, immunoprecipitation assays are typically performed in a fixed population of cells, so data represent an average cellular state. Thus, it is unclear if two modifications are actually present at a single site at the same time in individual cells. For example, it could be feasible that histone acetylation precedes RNAP2 phosphorylation, as we observed at the tandem gene array^22^. In this case, the two signals could be spatially anticorrelated at any given time, i.e., they would be spatially segregated.

To test if our data supported either of these two scenarios, we generated scatterplots from the signal intensities found within the nuclear pixels of all imaged cells. Here, intensity is rescaled from -1 to 1, with *I*_RS_ = (*I* − ⟨*I*)/(*I*_9.75_ − *I*_2.5_), where ⟨*I*⟩ is the average intensity and *I*_97.5_ and *I*_2.5_ are the 97.5 and 2.5 intensity quantiles, respectively (**Fig. 1C**). This analysis revealed a complex relationship between the various signals. For example, while the signals were overall positively correlated, especially the H3K27ac and H2B signals (**Fig. 1C, right**), the correlations were not linear. In particular, in the H3K27ac versus RNAP2-Ser5ph scatterplot (**Fig. 1C, left**), we observed a negative correlation in the upper-right quadrant, where both signals are relatively enriched. The H2B and RNAP2-Ser5ph had a strong negative correlation, consistent with dechromatization being coupled to transcription activity (**Fig. 1C, middle**). Thus, there is a mixture of distinct correlations between signals within our dataset, some positive and some negative. This would suggest the relationship between chromatin, histone acetylation, and RNAP2 phosphorylation is heterogenous and most likely context dependent.

### Tracking histone acetylation and RNAP2 phosphorylation dynamics at thousands of endogenous chromatin sites reveals two distinct dynamics

To better assess the complex relationships we saw emerging in our scatterplots, we next zoomed in on individual chromatin sites to track signals at those sites through time. To achieve this, we created an image processing pipeline consisting of 10 steps (**Sup. Fig. 2A**). Briefly, after averaging over five frames and correcting for cell movement (**Sup. Fig. 2B**,**C**), we used a local adaptive binarization filter to mask, identify, and track thousands of individual chromatin sites that were enriched with either H3K27ac or RNAP2-Ser5ph (**Sup. Fig. 2D-F**). An example is shown in **Fig. 2A**, where two individual sites (of many) from a sample cell are tracked through time, the first enriched with H3K27ac (green) and the second enriched with RNAP2-Ser5ph (purple).

**Fig. 2:**
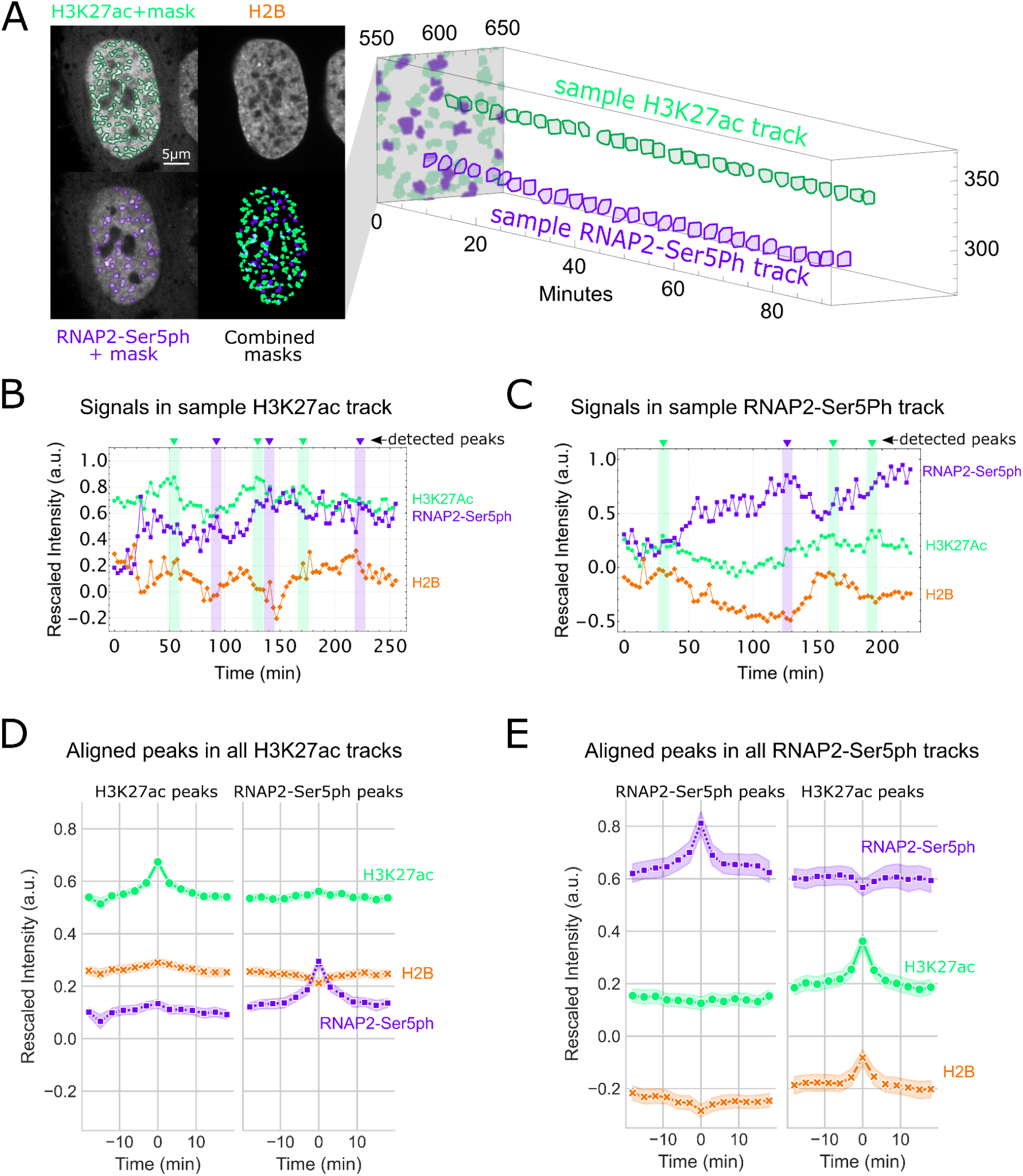
Hours-long tracking of chromatin regions enriched for H3K27ac and RNAP2-Ser5ph. Left) A local adaptive binarization filter is used to identify and isolate regions of chromatin enriched for specific post-translational modifications. Right) Enriched regions are tracked through time. An individual sample of H3K27ac enriched and RNAP2-Ser5ph enriched tracks are shown. **B.** Time traces from a sample track of chromatin enriched for H3K27ac, with average levels of H3K27ac, RNAP2-Ser5ph, and H2B within the region plotted. Identified peaks within H3K27ac and RNAP2-Ser5ph signals are labeled in green and purple, respectively. **C.** Same as B, for RNAP2-Ser5ph enriched region. **D.** Left) Aligned H3K27ac (n=2228 peaks) peaks from all H3K27ac enriched chromatin tracks (n=10,290 tracks total; n = 2743 greater than 30 frames). All signals were aligned based upon the detected peak’s timepoints. Right) Aligned RNAP2-Ser5ph detected peaks (n=2387) in the same tracks.

In total, we tracked 15,295 non-specific endogenous chromatin sites spread across the nuclei of 27 cells, 10,290 enriched with H3K27ac and 5,005 enriched with RNAP2-Ser5ph (**Sup. Fig. 3**). The sites varied in size, with a median cross sectional area of 1.12 ± 0.96 μm² (SD) for HK27ac-enriched sites and 0.92 ± 0.70 μm² (SD) for RNAP2-Ser5ph-enriched sites (**Sup. Fig. 3C**). These domains were too big to be single TADs^59,60^ or clutches^19^, but rather more on the order of chromatin territories^61^. Consistent with the negative correlation we saw in the upper quadrant of our RNAP2-Ser5Ph vs H3K27ac scatterplot, the two sites tended to border one another but were practically independent, with an overlap that occupied ∼5% of the nucleus or less (see, for example, the combined masks and quantification in **Sup. Fig. 3A**,**B**). In other words, the data confirmed regions of chromatin most enriched with H3K27ac had only moderate amounts of RNAP2-Ser5ph, and vice versa (**Sup. Fig. 3D**).

Given the spatially independent nature of chromatin sites enriched with H3K27ac or RNAP2-Ser5ph, we next wondered if the two types of sites displayed different dynamics. To address this question, we quantified the rescaled fluorescence intensities of each track through time (**Fig. 2B**,**C**). Analyzing these tracks in detail revealed a heterogeneous track-to-track population with a number of interesting behaviors and trends. First, many regions of chromatin maintained their enriched status (in either H3K27ac or RNAP2-Ser5ph) for half an hour or longer, with only minor fluctuations (**Sup. Fig. 2C, bottom**). When fluctuations did occur inside of H3K27ac enriched areas, we noticed that all three signals tended to shift in sync (**Fig. 2B**). Conversely, inside of RNAP2-Ser5ph enriched chromatin, increases in transcription signal tended to be accompanied by decreases in the H3K27ac and H2B signals and vice versa (**Fig. 2C**).

To assess the generalizability of these trends, we aligned all H3K27ac and RNAP2-Ser5ph signal peaks in both sets of tracks (see small arrows in **Fig. 2B**,**C** for a few sample peaks). We then examined the behavior of all signals when one signal peaked. For example, how H2B and RNAP2-Ser5ph responded when H3K27ac reached a local maximum. As expected, this revealed the dynamics of H3K27ac and RNAP-Ser5ph differed in the two types of sites. In H3K27ac-enriched sites, local peaks in one modification were predictive of small local peaks in the other modification, whereas in RNAP2-Ser5ph enriched sites, local peaks in one modification were predictive of small local troughs in the other modification (**Fig. 2D**,**E**). This opposing behavior, while subtle, was consistent across experimental replicates (**Sup. Fig. 4A**,**B**) and independent of whether we aligned signals by peaks (**Fig. 2D**,**E, Sup. Fig 4A**,**B**) or troughs (**Sup. Fig. 4A**,**B**). To further confirm the opposing behavior, we calculated the cross-correlation between the two modifications using signals from all tracks at all timepoints. In line with our peak alignment results, this demonstrated the two modifications are positively correlated with one another in sites enriched with H3K27ac but negatively correlated with one another in sites enriched with RNAP2-Ser5ph (**Sup. Fig. 4C**).

Left) Aligned RNAP2-Ser5ph peaks (n=874 peaks) from all RNAP2-Ser5ph enriched chromatin tracks (n=5,005 tracks total; n = 965 greater than 30 frames). All signals were aligned based upon the detected peak’s timepoints. Right) Aligned H3K27ac peaks (n=683) in the same tracks.Based on these observations we can conclude three things. First, the fact that peaks did not always align with peaks (and troughs did not always align with troughs) confirmed our analysis was free of focus drift or illumination issues that would cause signals to artifactually dim or brighten in sync. Second, despite H2B and H3K27ac being present together within nucleosomes, our peak analysis found the two signals can be decoupled when acetylation levels are high. Specifically, when we aligned RNAP2-Ser5ph peaks occurring inside highly acetylated tracks, a distinct trough in H2B appeared despite an increase in H3K27ac (**Fig. 3D, right**). Conversely, when we aligned RNAP2-Ser5ph troughs in the same tracks, a distinct peak in H2B appeared despite a drop in H3K27ac (**Sup. Fig. 4A, bottom-right**). These data therefore provide direct evidence that H3K27ac can act independently of and even counter changes in nucleosome density. Third, the temporal correlation we measured between H3K27ac and RNAP-Ser5ph dramatically changed depending on which modification was enriched. When H3K27ac was enriched, the signals fluctuated in a correlated manner, whereas when RNAP2-Ser5ph was enriched, the signals fluctuated in an anticorrelated manner (**Sup. Fig. 4C**). We therefore conclude chromatin enriched with either H3K27ac or RNAP2-Ser5ph display distinct dynamics from one another on the minutes timescale.

**Fig. 3:**
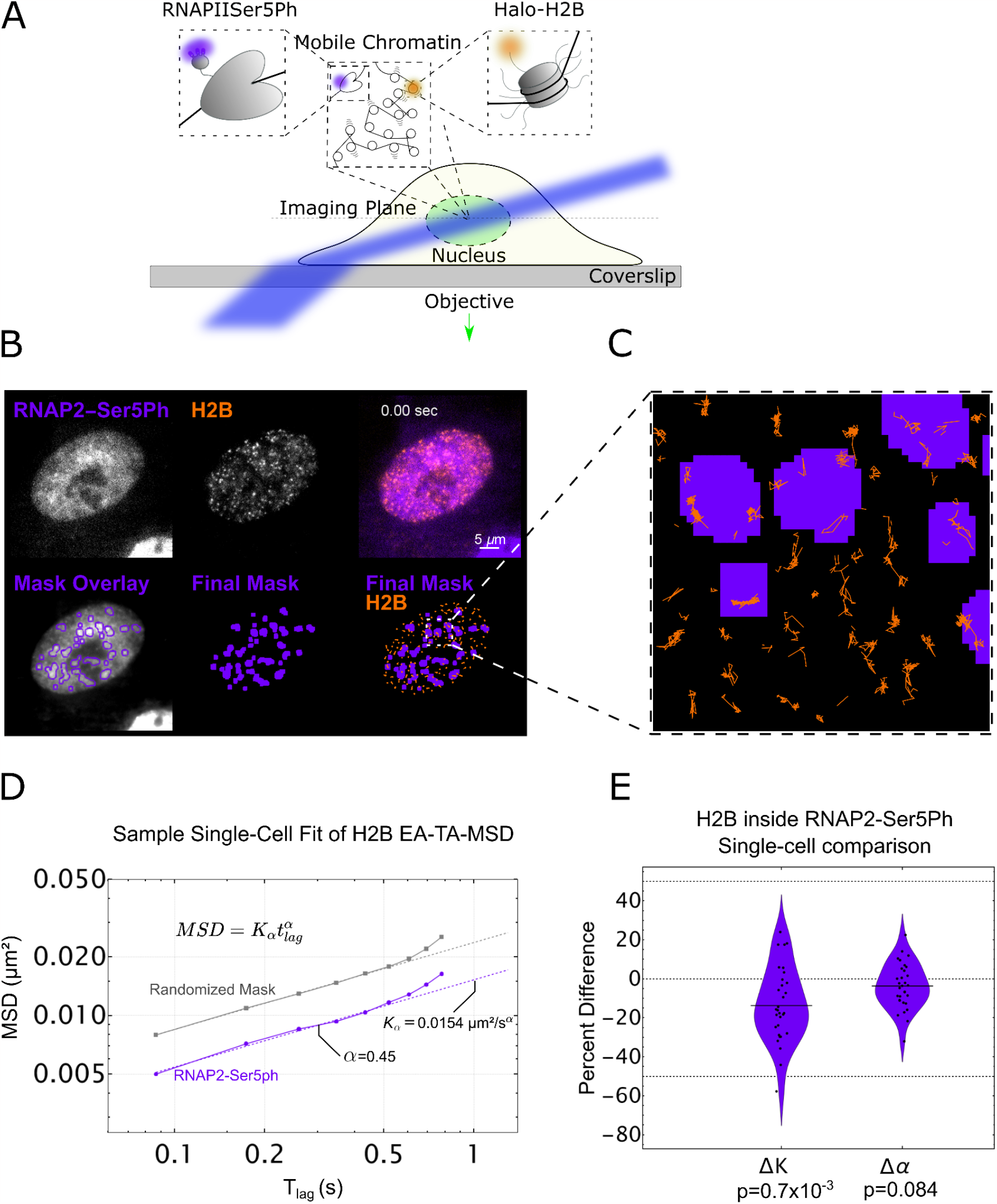
Single-particle tracking of H2B located within transcriptionally active regions marked by RNAP2-Ser5ph. **A.** Imaging scheme for single-particle H2B tracking experiments utilizing HILO microscopy setup. A combination of H2B-Halo and Fab specific for RNAP2-Ser5ph are utilized to track single molecules of H2B in the context of transcription. **B.** Top Row: Left) Sample single frame of RNAP2-Ser5ph Fab in living RPE1 cells. Middle) Sample single frame of single-molecule H2B-Halo in the same cell. Right) Colored composite of RNAP2-Ser5ph and H2B channels in purple and orange, respectively. Bottom Row: Left) Outlines of RNAP2-Ser5ph local adaptive binarization mask overlaid upon the 100-frame average image from which the mask was extracted. Middle) Single local adaptive binarization mask of RNAP2-Ser5ph enriched regions. Right) RNAP2-Ser5ph mask with overlaid H2B-Halo tracks in orange. **C.** Selected zoom from (B) showing sample full traces of Halo-H2B tracks occurring both inside and outside RNAP2-Ser5ph enriched regions. **D.** Sample single-cell fit for the ensemble MSD of all H2B tracks lasting 10 frames or longer, localized in a randomized mask control (gray, n=5860 total tracks across 10 randomizations) or in RNAP2-Ser5ph enriched, transcriptionally active regions (purple, n=463 tracks). Determined diffusion coefficient (K_α_) and alpha coefficient (α) for this fit are labeled.

### Single Particle Tracking of H2B at Transcriptionally Enriched Sites Reveals Chromatin Slowdown

Given the differences in the temporal correlations we observed between sites enriched with H3K27ac and RNAP2-Ser5ph, we were curious if the two types of sites also exhibited different dynamics on shorter length and timescales. In particular, we wondered if chromatin within each site exhibited unique microscopic dynamics that could help explain or predict our longer-term observations. Our experimental system was uniquely poised to address this question because we could adjust our imaging setup in a straight-foward manner to enable the tracking of individual nucleosomes on the milliseconds time scale while still co-imaging PTMs using FabLEM (**Sup. Fig. 1C**,**D**), and single-particle tracking and the corresponding analysis of trajectories is a useful way to decode the dynamics of intracellular components^62–65^.

We first focused on transcriptionally active regions marked by RNAP2-Ser5ph. We hypothesized nucleosomes in these regions would either diffuse faster or slower than normal. In support of slower nucleosomes, several recent reports^33,38,66^ have shown an anti-correlative relationship between chromatin mobility and transcriptional activity, consistent with the presence of hypothesized transcription factories or hubs that lock down chromatin^44^. In support of faster nucleosomes, on the other hand, it has been observed that some enhancers and promoters are more mobile after differentiation-induced transcription activation in embryonic stem cells, leading to the proposition that RNAP2 transcription activity stirs the local chromatin environment^52^.

To see which of these two opposing scenarios is dominant, we again bead-loaded RPE1 cells stably expressing H2B-Halo with AF488 labeled Fab specific to RNAP2-Ser5ph. To enhance fluorescence signal-to-noise and enable long-term tracking of single fluorophores, we switched from confocal to highly inclined laminated optical (HILO) sheet microscopy^36^ (**Fig. 3A**). To acquire many tracks with minimal crossing events, we sparsely labeled H2B-Halo with a reduced form of TMR ligand that is stochastically photoactivated during imaging^67^. Finally, we adjusted our imaging rate to a little over ten frames per second to exclusively track chromatin-incorporated H2B, an indicator for ‘single nucleosomes’^35,38^ (**Fig. 3B**).

Using this experimental setup, we collected tens of thousands of tracks from 31 cells across 3 experimental replicates and created an analysis pipeline (**Sup. Fig. 5A**). To isolate tracks associated with RNAP2-Ser5ph, we again applied a local adaptive binarization filter to the nuclei of each cell to highlight all sub-nuclear regions enriched with the specific PTM **(Fig. 3B, Sup. Fig. 5B**,**C**). We then selected all tracks that remained inside of the mask for at least ten consecutive frames (**Fig. 3C, Sup. Fig. 5D**). This left us with hundreds to thousands of single-nucleosome tracks per cell to quantify chromatin mobility in the context of RNAP2-Ser5ph.

To ensure that our masking process was not creating a selection bias amongst our tracks, we created a control mask. For this, we isolated and identified each region of connected pixels in each mask, and each region was then separately moved by a randomized vector. This procedure created a random mask with a similar morphology to our original mask, but randomly dispersed throughout the nucleus (**Sup. Fig. 5E**,**F**). In so doing, we could compare nucleosome dynamics inside the original RNAP2-Ser5ph mask to average nucleosome dynamics inside a large collection of morphologically identical random masks. This process is critical because restricting tracks to any masked region inherently selects for slower tracks and the amplitude of this selection bias depends on the precise size, shape, and morphology of the mask in question (**Sup. Fig. 5G**).

To quantitatively compare dynamics in our original and random masks, we fit the ensemble and time-averaged mean-squared displacement (MSD) of nucleosomes to a model of anomalous diffusion^68,69^ such that 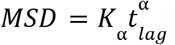. Here, *K*_⍺_ is the so-called generalized diffusion coefficient and *α* is the anomalous exponent (**Fig. 3D**). Similar to other reports^35,38,70^, our fitted *K*_*α*_ was in the range of 0.03 ± 0.01 μm^2^/sec, while our fitted *α* was consistently ∼0.45 ± 0.1 (even after correcting for both static and dynamic localization errors and eliminating their effect using a resampling approach^71^). Normal diffusion would be characterized by α=1. In our case the exponent α indicates the nucleosomes in the cell nucleus are subdiffusive, i.e., their MSD is sub-linear in lag time.

To control for cell-to-cell variability, we subtracted the fitted *K*_*α*_ and *α* values of nucleosomes inside randomized masks from those inside RNAP2-Ser5ph enriched masks. This generated a *Δα* and *ΔK* for each cell, allowing us to directly compare chromatin dynamics at a single-cell level and controlling for any cell-to-cell variability. As shown in **Fig. 3E**, the percent change in these differences provided a convenient single-cell measure of the impact RNAP2-Ser5ph enrichment has on single-nucleosome mobilities.

Difference in K_α_ and α for RNAP2-Ser5ph enriched regions vs a randomized mask control. Each data point represents the difference in fit coefficients for a single cell fit (n=31 cells), relative to the mean value, with each cell comprising hundreds to thousands of H2B tracks. Significance determined via student’s t-test.According to our fits, nucleosomes diffusing near chromatin sites enriched with RNAP2-Ser5ph experienced no significant change in the type of diffusion they underwent (*Δα ∼ 0*, **Fig. 3E, right**). Despite that, the nucleosome diffused ∼15% more slowly, characterized by a marked decrease in the diffusion coefficient. This substantial reduction in the mobility of nucleosomes near sites of transcription initiation supports the transcription factory/hub model^44^. Furthermore, because our measurements were at random sites in a random set of cells, our data suggest nucleosome slowdown near transcription initiation is a common event that does not depend on specific perturbations or drug treatments.

### Enrichment of H3K27ac Predicts an Increased Nucleosome Diffusion

We next turned our attention to histone acetylation, applying the same methodology to chromatin sites enriched with H3K27ac. According to the textbook model, histone acetylation should weaken DNA-nucleosome interactions, theoretically opening up chromatin structure to facilitate nucleosome mobility. In principle, we would therefore predict acetylated nucleosomes have a higher mobility than average. On the other hand, we just observed that less dense, transcriptionally active chromatin can have reduced mobility. The direct relationship between chromatin density and mobility is therefore not obvious. So far the only study^38^ that observed increased single-nucleosome mobility in the context of acetylation relied on the use of Trichostatin-A (TSA), a broad inhibitor of “HDAC” proteins that have many non-histone targets^72^. It is therefore unclear how chromatin will behave in regions enriched for H3K27ac.

To more directly address this question, we performed the same experiment as above, except we now loaded cells with AF488 labeled Fab specific for H3K27ac. As before, we tracked single nucleosomes, created real and randomized masks for H3K27ac enrichment, and isolated tracks inside of those regions (**Fig. 4A**). We then fit the ensemble and time-averaged MSD from each cell inside the masks to extract *K*_*α*_ and *α* (**Fig. 4B**), calculated *ΔK* and *Δα* for each cell, and plotted the distributions together (**Fig. 4C**).

**Fig. 4:**
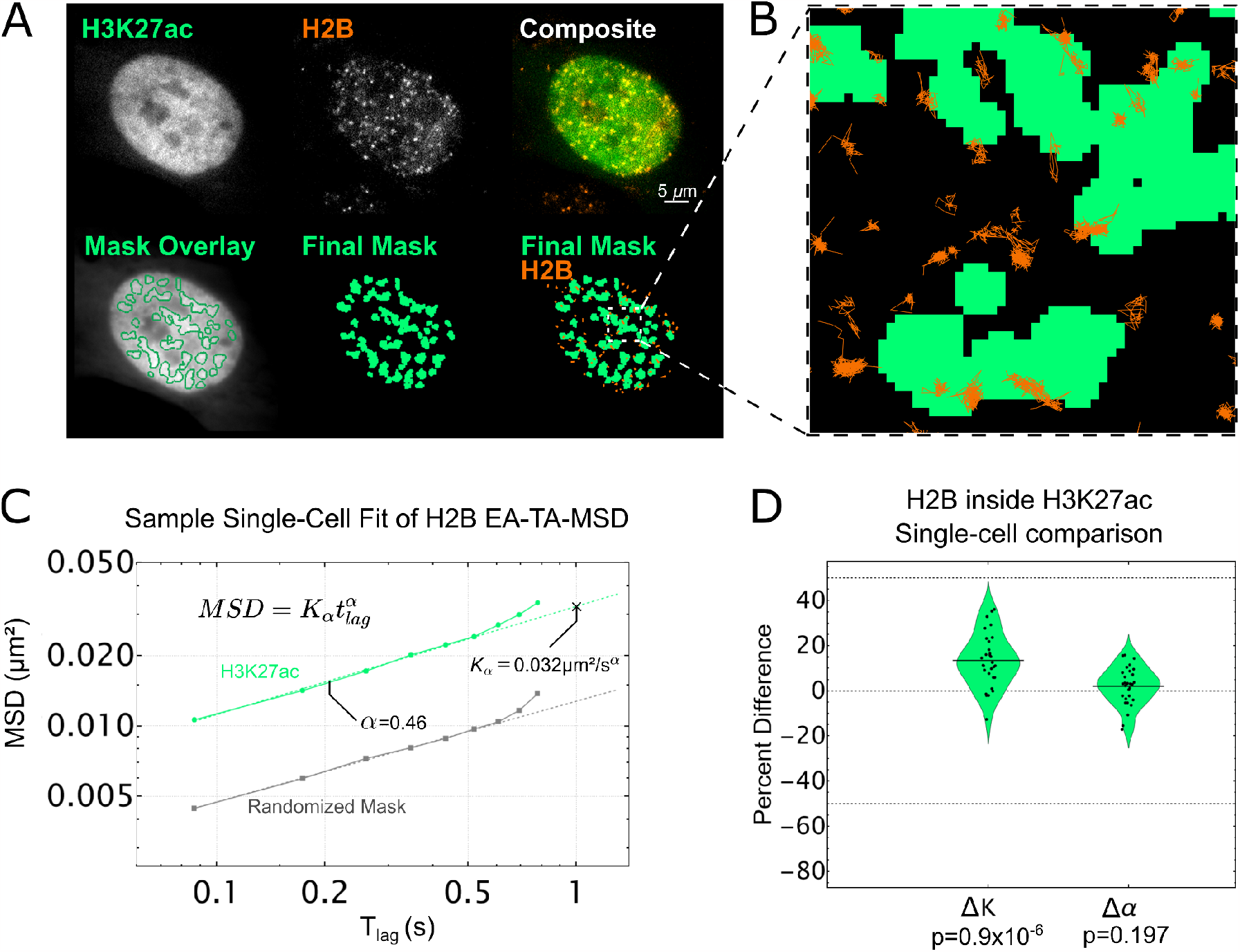
Single-particle tracking of H2B inside of H3K27ac enriched chromatin regions. **A.** Top Row: Left) Sample single frame of H3K27ac Fab in living RPE1 cells. Middle) Sample single frame of single-molecule Halo-H2B in the same cell. Right) Colored composite of H3K27ac and H2B channels in green and orange, respectively. Bottom Row: Left) Outlines of H3K27ac local adaptive binarization mask overlaid upon the 100-frame average image from which the mask was extracted. Middle) Single local adaptive binarization mask of H3K27ac enriched regions. Right) H3K27ac mask with overlaid Halo-H2B tracks in orange. **B.** Selected zoom from (A) showing sample full traces of Halo-H2B tracks occurring both inside and outside RNAP2-Ser5ph enriched regions. **C.** Sample single-cell fit for the ensemble MSD of all H2B tracks lasting 10 frames or longer, localized in a randomized mask control (gray, n=1892 total tracks across 10 randomizations) or in H3K27ac enriched chromatin (green, n=445 tracks). Determined diffusion coefficient (K_α_) and alpha coefficient (α) for this fit are labeled. **D.** Difference in K_α_ and α for RNAP2-Ser5ph enriched regions from a randomized mask control. Each data point represents the difference in fit coefficients for a single cell fit (n=30 cells), relative to the mean, with each cell comprising thousands of H2B tracks. Significance determined via student’s t-test.

As we saw with RNAP2-Ser5ph, our data indicate nucleosomes in regions enriched with H3K27ac did not experience any significant change in the type of diffusion they underwent (*Δα∼0*, **Fig 4D, right**). However, in stark contrast to what we saw with RNAP2-Ser5ph, we now observed a significant increase in the diffusion coefficient *K*_*α*_. Specifically, our data suggest nucleosomes diffuse ∼15% faster than normal when associated with chromatin enriched for H3K27ac (*ΔK>0*, **Fig 4D, left**). These data therefore provide direct live-cell support that acetylated nucleosomes move more freely in 4D space, as the standard model of histone acetylation would predict. Furthermore, the fact that we observed a speed up in nucleosomes in this case rather than a slow down suggests our general strategy for measuring nucleosome dynamics in the context of specific PTM masks is unbiased. Taken together, our data support a model whereby H3K27ac and RNAP2 Ser5ph mark opposing ends of the nucleosome mobility landscape, with H3K27ac marking sites of increased mobility and RNAP2-Ser5ph marking sites of decreased mobility.

## Discussion

Despite the well known correlation between histone acetylation, chromatin structure, and gene activity, the dynamic relationship between these factors has remained largely unexplored in living cells. In this study we addressed this issue by combining single-molecule tracking with live-cell imaging of PTMs. Using this unique combination of technologies, we quantified the dynamic interplay between endogenous histone acetylation marked by H3K27ac, endogenous transcription initiation marked by RNAP2-Ser5ph, and chromatin mobility marked by stably expressed H2B-Halo.

Our work revealed several fundamental relationships (**Fig. 5**). First, we showed chromatin that is highly enriched with H3K27ac was generally segregated from chromatin that was highly enriched with RNAP2-Ser5ph. This was a bit surprising given the strong and positive genome-wide correlation between H3K27ac and RNAP2-Ser5ph regularly observed via chromatin immunoprecipitation. A straightforward explanation for this apparent discrepancy is that individual chromatin sites oscillate between two states, one enriched with H3K27ac and one enriched with RNAP2-Ser5ph. Over time or across a cellular population this would result in an average co-enrichment in signals. Alternatively, given the apparent temporal stability of many of the regions we tracked, a single genetic locus could move back and forth between stable, preformed sites enriched with either H3K27ac or RNAP2-Ser5ph. Such models are not unprecedented as we observed a similar phenomenon at the MMTV tandem gene array^22^. In that study, H3K27ac levels at the array were high prior to activation, but the levels rapidly dropped post-activation when RNAP2-Ser5ph rapidly rose. Similarly, in another study with zebrafish embryos, high H3K27ac were observed at a developmental gene prior to its activation, but levels again dropped once RNAP2 was recruited^24^. Beyond these specific examples, it is now generally accepted that the vast majority of genes display bursty transcription^73–75^, transitioning between active and inactive states. In light of this, it is tempting to speculate that bursty genes also oscillate between PTM states, having high levels of H3K27ac at one time followed by high levels of RNAP2-Ser5ph at another time. Unfortunately, we were unable to confirm any state-transitions in this study because we could only track a specific site while it remained enriched. In the future this shortcoming could be reconciled by directly labeling and tracking single genetic elements while co-imaging H3K27ac and RNAP2-Ser5ph.

**Fig. 5:**
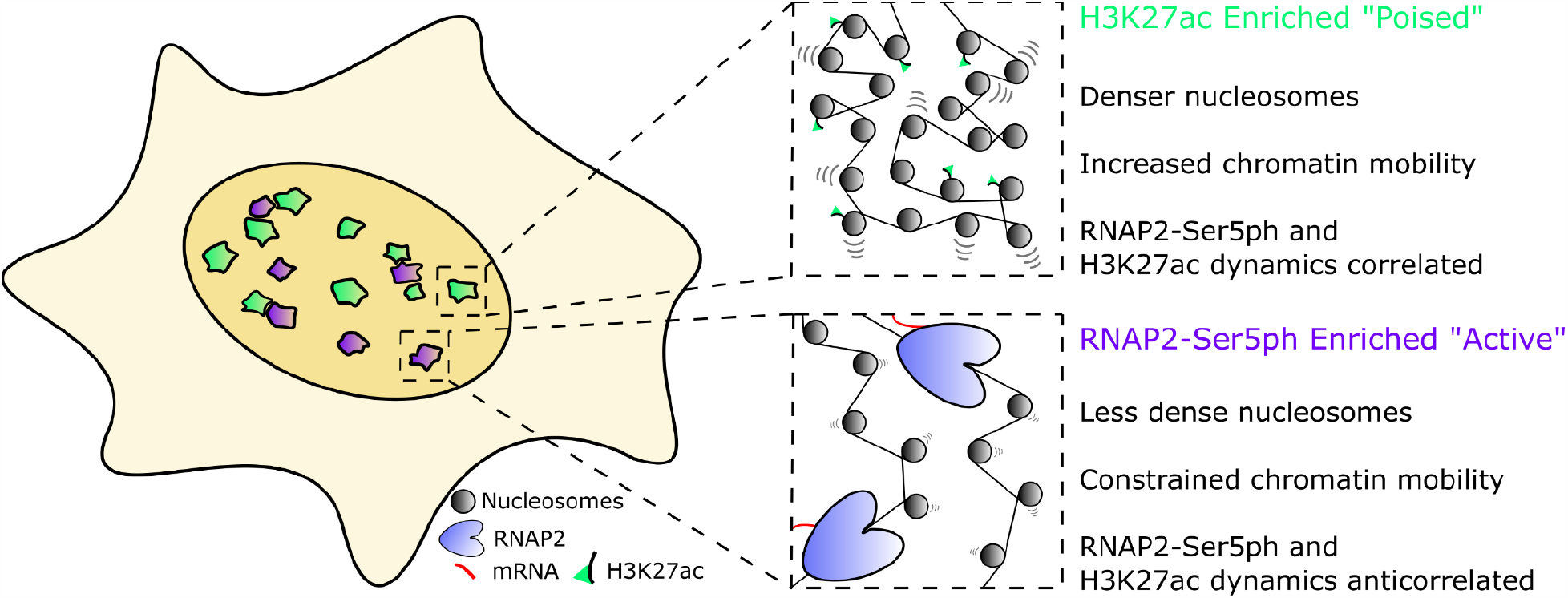
Two state model of RNAP2-Ser5ph and H3K27ac enriched chromatin domains. Inside the nucleus, two distinct sites are characterized by enrichment for RNAP2-Ser5ph or H3K27ac with little overlap. H3K27ac enriched regions represent poised chromatin, which is slightly more dense than average chromatin but has increased single nucleosome mobility. In combination, these factors reinforce 3D target searching, increasing the probability of contact between genetic elements or interacting proteins. RNAP2-Ser5ph enriched regions are areas of high transcriptional activity, where chromatin is decondensed but stabilized by the presence of bound transcription machinery.

Second, we showed the H3K27ac and RNAP2-Ser5ph signals were positively correlated through time in sites enriched with H3K27ac, but negatively correlated in sites enriched with RNAP2-Ser5ph. These distinct time-dependent spatial correlations persisted for up to 20 minutes (**Sup Fig. 4C)**, a timescale on par with the transcription cycle of average-sized genes according to RNAP2 FRAP and MS2-based gene tracking experiments^76–78^. The data therefore suggest the relationship between H3K27ac and RNAP2-Ser5ph may change throughout the transcription cycle. In sites ‘poised’ for transcription (which we define here as high H3K27ac and low RNAP2-Ser5ph), increases in H3K27ac levels help recruit RNAP2 and vice versa (green sites in **Fig. 5**). On the other hand, in transcriptionally ‘active’ sites (high RNAP2-Ser5ph, lower H3K27ac; purple sites in **Fig. 5**), increases in H3K27ac lead to a loss in RNAP2-Ser5ph and vice versa, most likely due to enhanced RNAP2 promoter escape or pause release by H3K27ac^13,22^ and the decondensation of nucleosomes by active RNAP2. This dual behavior is consistent with what we observed at the MMTV tandem gene array^22^. In that study, we showed H3K27ac at the array prior to its activation was predictive of (1) the efficiency of transcription factor recruitment immediately after hormone induction and (2) later RNAP2 promoter escape and chromatin decondensation. Whereas the former would positively correlate with RNAP2-Ser5ph, the latter would negatively correlate. Thus, our data support a model wherein H3K27ac plays multiple roles in tuning gene activation efficiency.

Third, we showed that single nucleosomes diffused ∼15% slower than normal in transcriptionally active sites enriched with RNAP2-Ser5ph and ∼15% faster than normal in sites enriched with H3K27ac. Taken together, these data paint a heterogenous picture of the chromatin mobility landscape^43^, one that is dynamically colored by local enrichments in specific PTMs (**Fig. 5**). While the molecular details of this two-state model have yet to be clarified, it does answer the conundrum between standard model predictions in acetylated and transcriptionally active regions. Namely, H3K27ac and RNAP2 can have separate impacts on chromatin landscape and mobility because they are acting in different places and times in living cells.

Regarding the slowdown of nucleosomes we observed in the context of transcription, our results are consistent with previous reports tracking single-genomic loci^33,66^, as well as with a similar study tracking single nucleosomes in the presence of transcriptional inhibitors^38^. Going beyond those previous studies we have now linked the slowdown to RNAP2-Ser5ph, a specific PTM associated with transcription initiation/pausing^7,14^. Furthermore, we did so without the use of drugs or broad-acting perturbations. Our data therefore support a model wherein transcription initiation is processed predominantly in factories^44^ or hubs^50,51^ that tend to lockdown nearby chromatin rather than stir it up^52^. While the lockdown we observe is consistent with traditional notions of transcription factories, we cannot rule out other models of lockdown. For example, crowding within phase-separated condensates^79^ or within stiff transcriptional loops^80^ could also restrict chromatin mobility. Similarly, we cannot exclude the possibility that some genetic loci behave differently. This is because we had no way of labeling the genomic context of each track. In other words, we could not say if a subset of fast or slow tracks were associated with a specific gene or a specific chromosomal locus. Furthermore, because we focused exclusively on RNAP2-Ser5ph, we may have missed some portion of transcription sites. Recently, there is growing evidence that transcription initiation (marked by RNAP2-Ser5ph) and elongation (marked by RNAP2-Ser2ph) are spatially separated^12,81^. Thus, it is possible that chromatin enriched with actively elongating RNAP2 may still associate with highly mobile nucleosomes^81^. In the future it will therefore be interesting to compare nucleosome mobilities within the context of a wider range of RNAP2 PTMs.

Regarding the enhanced mobility of nucleosomes we observed in the context of H3K27ac, our results are again consistent with a previous report that tracked nucleosomes in the presence of inhibitors^38^. As before, we go beyond this work by directly tracking nucleosomes in the context of a specific PTM without the use of inhibitors, which have many off-target and non-specific effects. Furthermore, by focusing on the H3K27ac modification, a well known marker of promoters and enhancers^55^, our data suggest these elements are especially dynamic prior to transcription activation, in line with the predictions of the standard model of gene activation by histone acetylation. Although we did not examine other residue-specific histone acetylation marks, such as H3K18ac or H3K9ac, we expect many will behave similarly, not only because of the earlier experiments with TSA^38^, but also from the classic viewpoint that acetylation acts redundantly to neutralize histone charge^3^. In particular, we expect H3K27ac is most likely redundant because it is written by the promiscuous p300/CBP lysine acetylase subfamily^82^ and its complete removal has little impact on fly development and mouse embryonic stem cell differentiation^83,84^. Nevertheless, it will be interesting to examine other acetylations one by one. H3K9, in particular, would be an interesting comparison since it plays a distinct role in nuclear receptor transactivation and is written independently of H3K27ac by the Gcn5/PCAF lysine acetylase subfamily^85^.

In summary, we have combined single-nucleosome tracking with live-cell, multicolor PTM imaging to reveal two types of distinct intranuclear regions that are enriched with RNAP2-Ser5ph or H3K27ac and that display distinct dynamics across multiple time and length scales. We believe this imaging strategy, combined with computational tools to track, extract, and analyze dynamics within masked regions of cells in an unbiased fashion, create a powerful experimental platform to better elucidate the role residue-specific PTMs play in gene expression networks. The techniques we have developed here can now be used to study a suite of other modifications. For example, beyond acetylation, it will be interesting to mask chromatin marked by other histone PTMs, both individually and in combination. Our methodology to track nucleosomes in the context of PTMs can also be generalized to track other proteins, such as transcription factors or chromatin remodelers. As the number of PTM-protein combinations is huge, our general method can be broadly applied to help elucidate the hidden roles PTMs play in diverse gene and chromatin regulatory networks.

## Acknowledgements

The authors thank all members of Stasevich lab, past and present, for their discussion and insight. We also thank Jeffrey Hansen for his helpful discussions and advice. We also thank Luke Lavis for the generous gift of JFX-554. This work was supported by a grant from the National Institutes of Health (R35GM119728) to T.J.S.

## Contributions

Conceptualization: M.S. and T.J.S. Performed experiments / collected data: M.S. and T.M. Antibodies and Fab Preparation: M.S. and H.K. Cell Culture: M.S. Software development and implementation: M.S., D.K., T.M., T.J.S. Diffusion Analysis: M.S. and D.K. Original draft: M.S. and T.J.S. Review and edits of drafts: M.S., T.M., D.K., H.K. and T.J.S. Resources, supervision and funding: T.J.S.

## Ethics Declarations

The authors declare no competing interests.

## Materials And Methods

### Cell culture and care

Human retinal epithelial cells (hTERT-RPE1, ATCC) were grown in 10% (v/v) FBS (Atlas) / DMEM (Thermo Scientific). Media was supplemented with 1 mM L-glutamine (Gibco) and 1% (v/v) Pen/Strep (Invitrogen/Gibco) and cells were grown at 37 °C in 5% CO2. Cell density was maintained between 15–80%. RPE1 cells were purchased from ATCC and were authenticated via STR profiling by ATCC and tested negative for mycoplasma contamination. Cells were stored at -80° C in Cellbanker1 (Amsbio LLC), and thawed >1 week prior to any imaging.

### Stable cell line creation

RPE1 cells were transfected with a Halo-H2B plasmid containing a Neomycin-resistance selection site. Transfection was performed with an LTX Lipofectamine with Plus Reagent kit (Thermo Fisher), per the manufacturer’s instructions. Briefly, an 80% confluency, 35mm MatTek Chamber was washed and the media was replaced with 1.75 mL Opti-MEM (Thermo Scientific) directly before transfection. The transfection solution included 2.5 μg DNA plasmid, 7.5 μL Plus reagent, and 7.5 μL Lipofectamine, and the remainder Opti-MEM for a total solution volume of 250 μL. This solution was incubated for 5–15 min at room temperature before being added to the cell chamber. Cells were incubated in this transfection solution for 2–4 h before the media was changed back to 10% FBS-DMEM. Cells were then transferred to a 10cm dish and allowed to replicate for 14 days under selection at 750μg/ml Geneticin G418 Sulfate (Gibco). Cells were then stained with Halo-JF646 at 500nM for 30 min, immediately washed with 1X PBS, and sorted via flow cytometry (FACS) to obtain a consistent population of stable H2B-Halo expressing RPE1 cells. The stable line was maintained under a constant selection of 400μg/ml Geneticin.

### Fab generation and dye conjugation

RNAP2-Ser5Ph and H3K27ac specific Fab were generated and affinity purified as previously described^21^. Briefly, Fab were generated from monoclonal antibodies using the Pierce Mouse IgG1 Fab and F(ab’)2 Preparation Kit (Thermo Fisher). Antibodies were first digested into Fab in a Zeba Desalt Spin Column (Thermo Fisher) containing immobilized Ficin while gently rotating for 3–5 h at 37°C. Fab were purified from the digest by centrifugation in a NAb Protein A column. Eluted Fab were concentrated to >1 mg/mL and stored at 4°C. Fab labeling with AF488 and CF640 was performed in small batches using 100 µg Fab. The dye was an AlexaFluor488 ester (Invitrogen) or CF640 (Biotium) dissolved in DMSO and either used immediately or stored at −20 °C. For labeling, 100 µg of Fab was dissolved in a final volume of 100 μL of 100 mM NaHCO_3_ (pH 8.5) plus 4 µl of AF488 or 2 µl of CF640 dye. Fabs were incubated with dye for ∼2 h at room temperature with constant gentle rotation and agitation. The Fab were separated from unconjugated dye in an equilibrated PD MiniTrap G-25 desalting column (GE Healthcare). Fab were concentrated in an Amicon Ultra-0.5 Centrifugal Filter Unit (NMWL 10 kDa; Millipore) to >1 mg/ml. The degree of labeling (DOL) was calculated using the following equation:

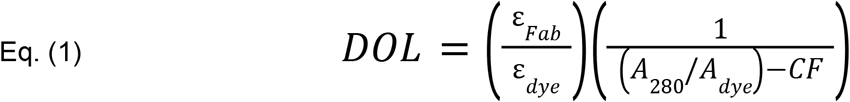

ε_Fab_ is the extinction coefficient of Fab (70,000 M^−1^ cm^−1^), ε_dye_ is the extinction coefficient of the dye used for conjugation (71,000 M^−1^ cm^−1^ for AF488 and 105,00 M^−1^ cm^−1^ for CF640), A280 and Adye are the measured absorbances of dye-conjugated Fab fragments at 280 nm and at the peak of the emission spectrum of the dye (488 nm for AF488, 637 nm for CF640), respectively, and CF is the correction factor of the dye (the ratio of the absorbances of the dye alone at 280 nm to at the peak). If the DOL was <0.7, this protocol was repeated on the same Fab to increase their DOL to ∼1. Fab were then stored at 4°C for later use.

### Bead loading of Fab and Halo-ligand staining prior to imaging

To bead load RNAP2-Ser5Ph and H3K27ac specific Fab, cells were plated at ∼75% confluency on a 35 mm glass-bottom chamber (MatTek) 24 hours before imaging. Cells were bead loaded as described previously^86^. Briefly, 1.5μg of each Fab required for an experiment is added together and diluted to 5μl in 1x PBS. Cell media is removed, and the 5 μL mixture of Fab was pipetted directly on top of the cells, followed by a sprinkling of ∼100 μm glass beads to form a monolayer on top of the cells (Sigma Aldrich). The entire chamber is then lifted and firmly tapped on the worktop 6-8 times inside of the culture hood. The media was then immediately replaced and cells were returned to the incubator. After ∼4 h of recovery, cells were washed three times in phenol-free DMEM with 10% FBS and 1 mM L-glutamine. For single molecule experiments, HaloTag-TMR ligand was first incubated in 100mM NaBH_4_ / 1X PBS for 10 minutes, and then diluted to 500pM and cells were stained. For non-single molecule experiments, HaloTag-JaneliaFlour-X554 (JFX554) was used at 200nM concentration. In all cases cells were incubated with HaloTag ligand for 30 minutes, then followed by 3x (3 washes in DMEM with 10% FBS and 1 mM L-glutamine followed by 5 minutes of incubation). Cells were moved to the microscope stage-top incubator for imaging ∼4h post-bead loading.

### Live cell imaging with the confocal microscope

Live cell images were acquired on an Olympus (IX83) inverted spinning disk confocal microscope equipped with a Cascade II EMCCD camera. A 100x oil immersion objective with a pixel size of 0.096 μm was used for all images. Cells were plated 24 hours before imaging, beadloaded 4-5 hours before imaging, and stained with Halo-JFX554 30 minutes before imaging, as described above. The chamber was allowed to acclimate in the stage-top incubator for 2-3 hours before final image collection to prevent focal shift. 488 nm, 561 nm, and 637 nm lasers were used in all cases. 5 frames were acquired in each channel, for each cell, every 2 or 3 minutes.

### Confocal image registration

A sequence of post-processing functions was applied to all confocal images prior to any binarization or tracking of specific subnuclear regions. First, for each channel, the 5 images collected at each time point were averaged together. To correct for cell movement, a maximum intensity projection of all 100 time points of H2B images was created for each cell, and a mask of nucleus movement was drawn by hand around the maximum projection. The mask of nucleus movement was multiplied by all images, leaving only the nucleus of interest. The resultant nuclear H2B images were fed into the FIJI^87^ plugin Register Virtual Stack Slices^88^, which aligned each nuclear image. This created a list of transformation functions for each time point for each cell, which were then used in conjunction with FIJI plugin Transform Virtual Stack Slices to register the Fab channels for each cell, ensuring that the exact same transformation is applied to all channels.

### Identification of sub-nuclear enriched regions with local adaptive binarization

To isolate sub-nuclear regions enriched for RNAP2-Ser5Ph or H3K27ac, custom code was written which utilized a built-in Mathematica function, LocalAdaptiveBinarize. First, a nuclear mask was drawn by hand, and multiplied by all images to isolate only the nucleus. Each nuclear image had its intensities normalized from 0 to 1, and a function was applied to each pixel:

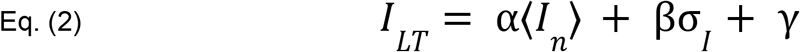

where *I*_*LT*_ is the local threshold, ⟨*I*_*n*_ ⟩ and *σ*_*I*_ represent the local mean and standard deviation in the neighborhood of *n* pixels. The values α, β, γ are user-defined constants. If a pixel’s value is higher than the resultant function, it is set to 1, else it is set to 0. For all masks created, the same parameters were used. These are: n=441 (21×21 window), α=0.94, β=0.6, γ=0.05. The resulting binary image had speckle noise eliminated by applying a 10-pixel size filter, and the remaining regions were dilated by 1 pixel to smooth and fill in gaps.

### Tracking of RNAP2-Ser5Ph and H3K27ac enriched regions

The local adaptive binarization masks from above were then tracked using FIJI plugin TrackMate^89^ (v7.6.1) using the following parameters: LoG Detector; Estimated Blob Diameter: 15.0; Pixel Threshold: 0.01; Sub-Pixel Localization: Enabled; Simple LAP Tracker; Linking Max Distance: 10 pixels; Gap-Closing Max Distance: 3 pixels, Gap-closing Max-Frame Gap: 2 frame. The resulting tracks were fed into custom Mathematica code which finds the region associated with each x-y coordinate for each track. A region was defined as the set of connected pixels with a value of 1 that are also within 5 pixels in any direction from the X-Y centroid. The values of each channel’s raw data pixels (H3K27ac, H2B, RNAP2-Ser5Ph) were then rescaled for each nuclear image, with -1 and 1 set to the 2.5 and 97.5 quantiles, respectively, to account for outliers. All 3 channels’ pixel intensities within the tracked region were then extracted and averaged together, resulting in a single value for H3K27ac, H2B, and RNAP2-Ser5Ph for each tracked region, for each time point. If the resulting time traces had missing data points due to track gaps, these values were linearly interpolated from adjacent data (less than 3% of total data).

### Correlation and Cross-Correlation Analysis

To calculate the correlation of H3K27ac, H2B, and RNAP2-Ser5Ph rescaled intensity traces, the Pearson correlation coefficient was calculated. The mean time-lag cross correlations were obtained by calculating the Pearson correlation coefficient for each track individually at various time lags, and then weighting each time lag by the length of the track used to calculate it. Then all tracks’ cross correlation curves were averaged. The error of cross correlation curves was obtained via bootstrapping. To bootstrap, intensity traces were randomly selected from the pool of data, with replacement, equal to the size of the dataset. Then, the mean cross correlation is calculated as above, and the process is repeated 1000 times. The error is reported as the standard deviation of all 1000 bootstrapped cross correlation curves.

### Peak and trough calling analysis

To find peaks in the H3K27ac and RNAP2-Ser5Ph rescaled intensity tracks, the scipy^90^ function in Python scipy.signal.find_peaks() was used using a ‘prominence’ of 0.15, a ‘width’ of 2 and ‘distance’ of 10. Troughs were found the same way after signals were inverted. For this analysis, tracks of length < 30 time points were ignored. Also, if peaks were within 10 time points of the beginning or end of a track, they were ignored (to facilitate easier alignment of all peaks).

### Live cell imaging on the HILO microscope

All live-cell imaging was performed on a custom built widefield fluorescence microscope with a highly inclined thin illumination scheme described previously. Briefly, the microscope equips three solid-state laser lines (488, 561, and 637 nm from Vortran) for excitation, an objective lens (60X, NA 1.49 oil immersion, Olympus), an emission image splitter (T660lpxr, ultra-flat imaging grade, Chroma), and two EMCCD cameras (iXon Ultra 888, Andor). Achromatic doublet lenses with 300 mm focal length (AC254-300-A-ML, Thorlabs) were used to focus images onto the camera chips instead of the regular 180 mm Olympus tube lens to satisfy Nyquist sampling (this lens combination produces 100X images with 130 nm/pixel). A single camera was used to capture both Fab (AF488, Thermo-Fisher) and Halo-H2B stained with Halo-ligand (TMR, Promega). A high-speed filter wheel (HS-625 HSFW TTL, Finger Lakes Instrumentation) is placed in front of the camera to minimize the bleed-through between the red and the green signals (593/46 nm BrightLine for the red and 510/42 nm BrightLine for the green, Semrock). All single molecule imaging was performed on a single focal plane for each cell. The laser emission, the camera integration, and the emission filter wheel position change were synchronized by an Arduino Mega board (Arduino). Image acquisition was performed using open source Micro-Manager software^91^ (1.4.22).

Live RPE1 cells were placed into a stage-top environmental chamber at 37 °C and 5% CO2 (Okolab) to equilibrate for at least 30 min before image acquisition. Imaging size and exposure time were set to 256 × 256 pixels and 30 msec, respectively. The resultant imaging rate was ∼23hz, (43.34 ms per frame). Fab and single-molecule images are alternated, resulting in 86.68 ms per frame for each channel. Laser powers were measured at the back focal plane to be 150 μW for 488 nm and 3.4 mW for 561 nm.

### Single-molecule H2B tracking and Mask Assignment

Single-molecule tracks were identified using TrackMate 5.0.1^92^ with the following parameters: LoG Detector; Estimated Blob Diameter: 5.0; Pixel Threshold: 100; Sub-Pixel Localization: Enabled; Simple LAP Tracker; Linking Max Distance: 3 pixels; Gap-Closing Max Distance: 2 pixels, Gap-closing Max-Frame Gap: 1 frame.

Movies were 5000 total frames, 2500 of Fab and 2500 of single molecule H2B in an alternating fashion. Masks were generated by first averaging 100 frames of Fab images, then applying the local adaptive binarization algorithm as above, thereby generating 25 masks which updated every ∼8.2 seconds. Single-molecule H2B tracks were then assigned to a mask based upon maximum residence time. Tracks were determined to be inside or outside of H3K27ac, RNAP2-Ser5Ph, or randomized masks based upon their sub-pixel X-Y coordinates. Tracks which crossed between mask borders were split, and track segments which resided in an area for at least 10 consecutive frames were kept.

### Diffusion analysis (Generating EA-TA-MSD curves, truncating tracks based on length, and fitting for K and alpha)

TrackMate files were fed into a custom MatLab script for analysis. To calculate the time-averaged mean square displacement (TAMSD) for each track, all tracks were first truncated to include only their first 10 frames. TAMSD was calculated as follows^93^:

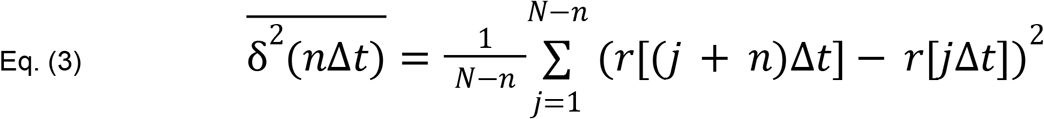

where *r* is xy position as a function of time, the overbar indicates a temporal average,∆*t* is the frame time, *t*_*lag*_ = *n*∆*t* is the lag time, and N is the number of data points in the trajectory. The ensemble averaged TAMSD (EA-TA-MSD) was calculated by averaging all single-nucleosome TAMSD results for each cell. A localization error of ∼30nm was roughly determined from the intercept of the EA-TA-MSD curve, and a correction was applied to all curves prior to fitting. The first five points of the logarithm of EA-TA-MSD vs logarithm of time curve for each cell were used in a linear fit to obtain *K* (generalized diffusion coefficient) and α (anomalous exponent) according to:

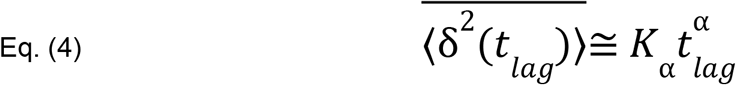

To obtain ∆*K*_α_ and ∆α for each individual cell, *K*_α_ and α obtained using randomized masks were subtracted from *K*_α_ and α obtained using the real-mask.

### Randomized mask generation

A randomized mask was created from the local adaptive binarization of real data through the following process. Each subnuclear component, defined as the series of connected foreground pixels within the local adaptive binarization, had its pixel locations identified. For each component, a randomized vector was created with value between -20 and 20 for both the x and y directions. Each component was then shifted in its entirety by pixels equal to the value of the randomized vector. The result was a randomized, binarized mask which has a morphological nature based on the data from which it was created, but with each subnuclear component randomly placed.

### Resampling MSD Curves to Calculate α WIthout Static and Dynamic Errors

MSD data obtained form single particle tracking are prone to static and dynamics errors, making it challenging to obtain a reliable estimation of the anomaly exponent ⍺^94,95^. In order to eliminate both types of errors, we employ a resampling approach as proposed by Weiss^71^. Namely, we analyzed nucleosome trajectories within a real and control mask with minimum lengths 20 or 30 frames. We employed two different trajectory lengths to ensure that the approach is robust in terms of number of frames. For each dataset, we resampled the trajectories, taking only even or odd positions, resulting in the two trajectories with frame times 2∆*t* for each original one with frame time ∆*t*. The TA-MSD was calculated for the original and resampled data, 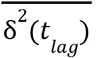. Then a translated MSD functional is computed for each 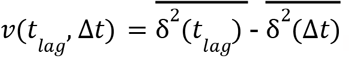, so that the exponent α is found from the relation:

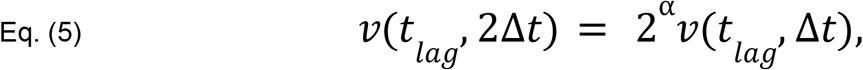

where *v*(*t*_*lag*_, 2∆*t*) is the functional for the resampled data. The exponent ⍺ is thus:

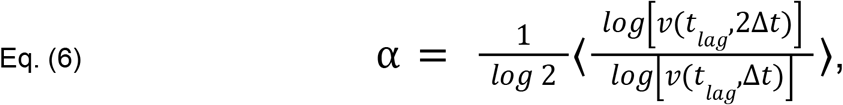

after averaging over different lag times. The α from three independent data sets were then averaged to find the anomaly exponent for H3K27ac and RNAP2-Ser5ph data, and for their associated control masks.

