## Supplementary Figures for "Live-cell imaging uncovers the relationship between histone acetylation, transcription initiation, and nucleosome mobility"

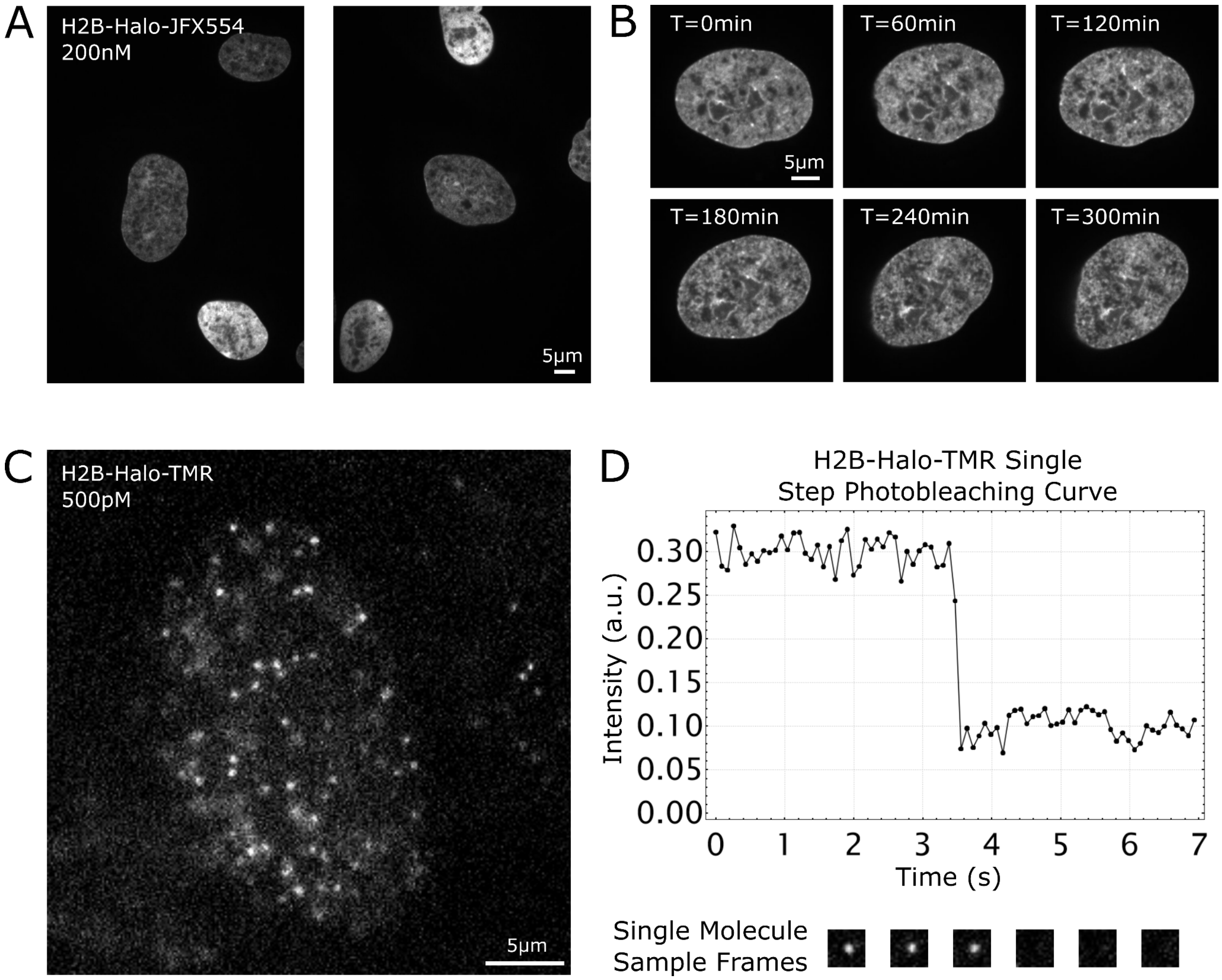


**Sup Fig. 1: H2B-Halo Stable RPE1 Cell Line Allows For Tracking Chromatin at Multiple Timescales**

A. Sample images of multiple RPE1 cells in a single frame, stably expressing H2B-Halo, stained with Halo-JFX554 dye at 200nM concentration.
B. Single cell H2B intensity and dynamics can be tracked for hours without significant photobleaching
C. Same RPE1 cells as in A), except stained with 500pM Halo-TMR. Now, single nucleosomes can be identified and tracked in real time with sub-second resolution.
D. Single-step photobleaching curve for a sample H2B track, along with frames of a tracked H2B before and after photobleach occurrence.

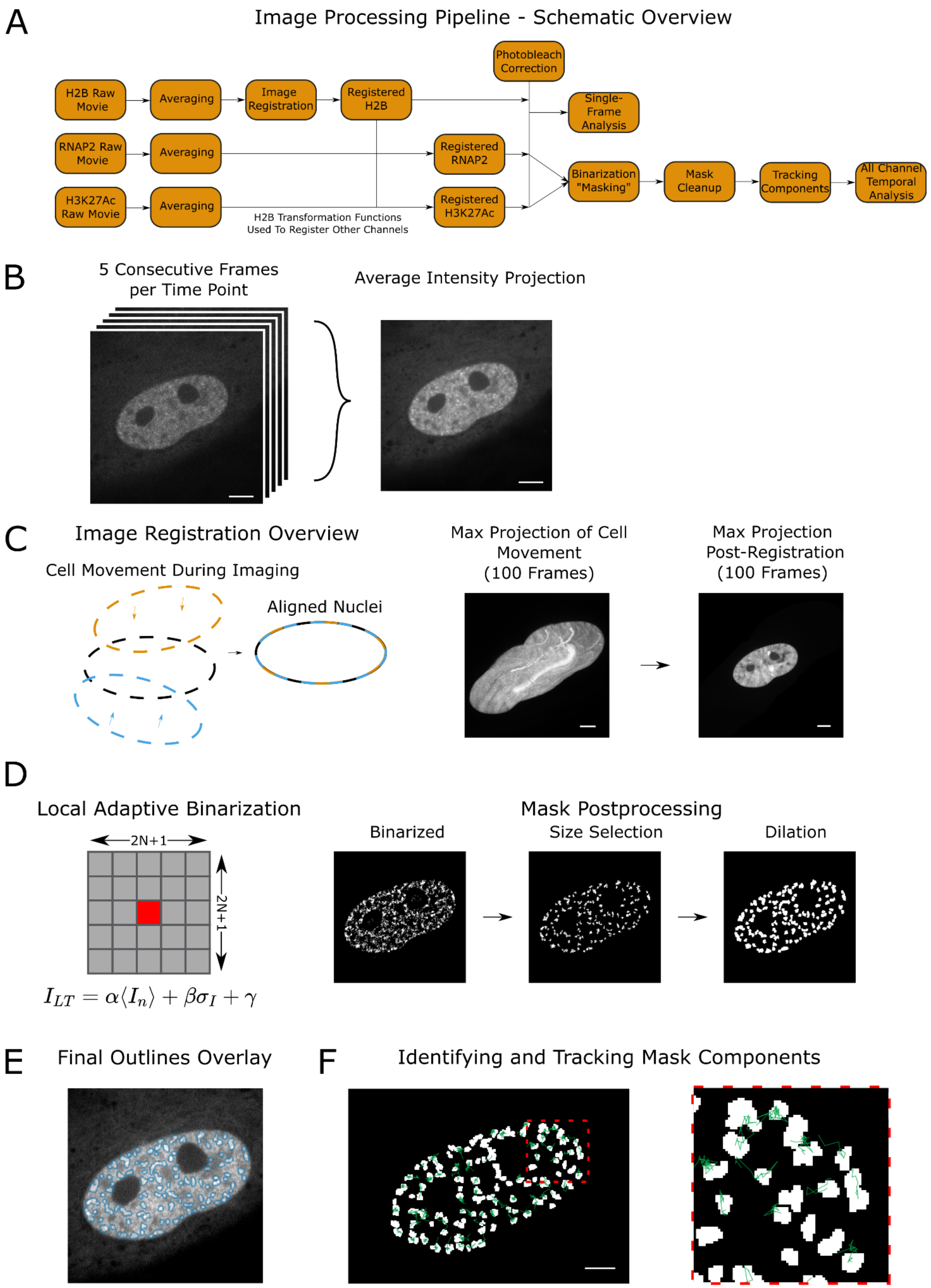
A
Image Processing Pipeline - Schematic Overview
H2B Raw Movie
Averaging
Image Registration
Registered H2B
Photobleach Correction
Single-Frame Analysis
RNAP2 Raw Movie
Averaging
Registered RNAP2
H3K27Ac Raw Movie
Averaging
Registered H3K27Ac
H2B Transformation Functions Used To Register Other Channels
Binarization "Masking"
Mask Cleanup
Tracking Components
All Channel Temporal Analysis
B
5 Consecutive Frames per Time Point
Average Intensity Projection
C
Image Registration Overview
Cell Movement During Imaging
Aligned Nuclei
Max Projection of Cell Movement (100 Frames)
Max Projection Post-Registration (100 Frames)
D
Local Adaptive Binarization
2N+1
2N+1
$I_{LT} = \alpha\langle I_n \rangle + \beta\sigma_I + \gamma$
Mask Postprocessing
Binarized
Size Selection
Dilation
E
Final Outlines Overlay
F
Identifying and Tracking Mask Components

**Sup Fig. 2: Analysis pipeline for confocal imaging of RNAP2-Ser5ph, H2B, and H3K27ac in living RPE1 cells**

A. Schematic overview of the analysis pipeline used to process confocal images and track regions of H3K27ac and RNAP2-Ser5ph.
B. Five frames are collected per time point and averaged to reduce noise from freely diffusing background Fab.
C. Each cell's nucleus is individually registered to the H2B channel to correct for any cell movement during imaging. Pre- and Post-registration max projections of an example cell are shown.
D. Local adaptive binarization is used to identify regions of a nucleus enriched for H3K27ac or RNAP2-Ser5ph. For additional information see materials and methods. The same binarization parameters are applied to all cells in all cases. After binarization, components less than 10 pixels are removed to reduce speckle noise, and then dilated by a single pixel to fill in gaps and smooth edges.
E. An outline of final binarization components (blue) overlaid on the original Fab image.
F. Demonstration of tracking individual H3K27ac chromatin regions through time. Components with a line trace (green) show the detected region's centroid movement over time. Components without a shown line trace did not meet the minimum requirement of 10 frame length for tracking and were discarded.

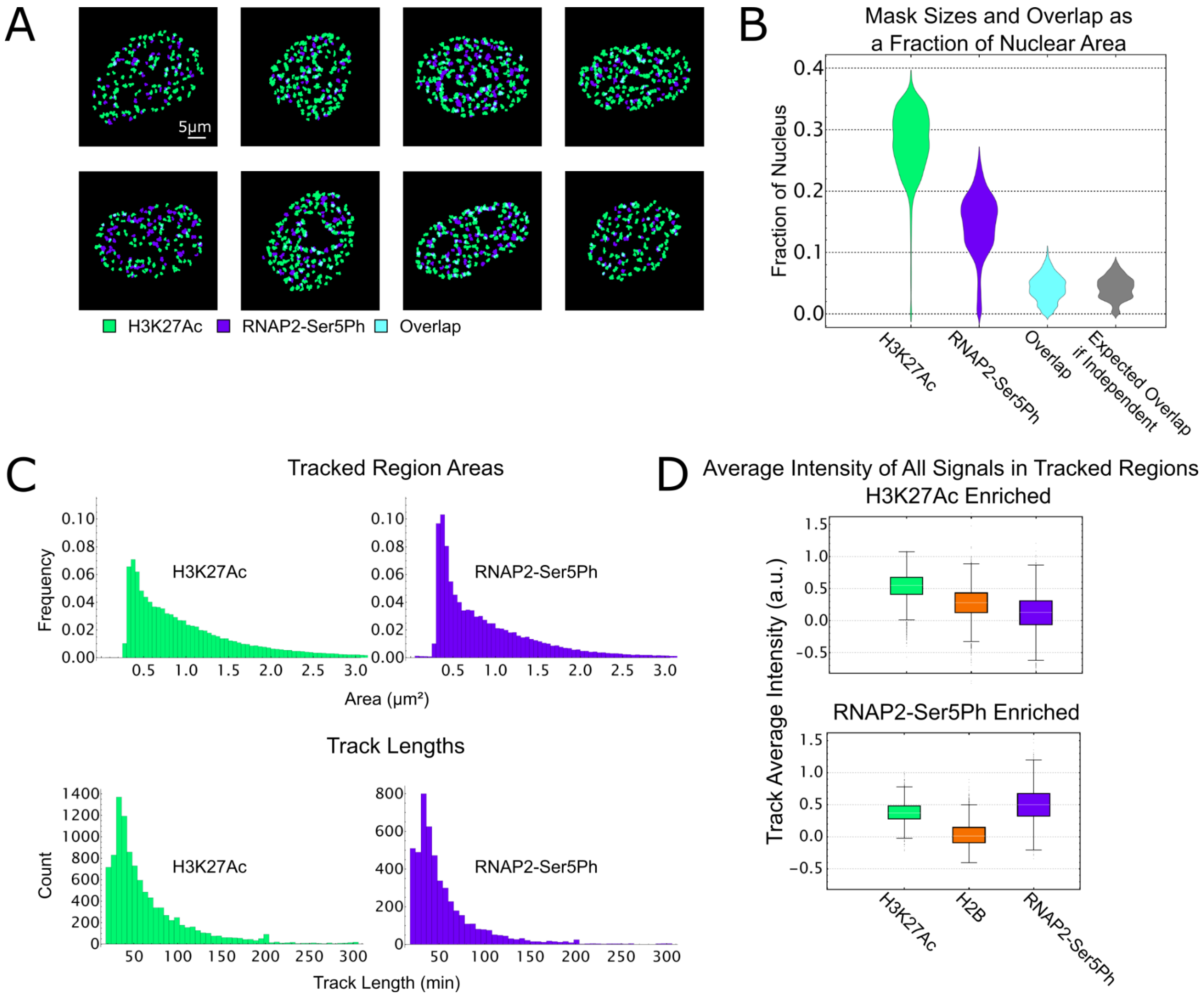


**Sup. Fig. 3: Characterization of H3K27ac and RNAP2-Ser5ph Regions: Overlap, Nuclear Area, and Intensity**

A. Sample images from 8 cells demonstrating the location and overlap between H3K27ac and RNAP2-Ser5ph masks.
B. Distribution of mask area as a fraction of nuclear area, with the overlap between masks also shown. Expected overlap if independent represents the amount of expected overlap if each mask's area were independent and uniformly distributed throughout the nucleus. n=27 cells, 2700 masks.
C. Top) Histogram of area of detected regions for both H3K27ac and RNAP2-Ser5ph masks. n=228,610 regions for H3K27ac, n=143,854 regions for RNAP2-Ser5ph. Bottom) Histogram of track lengths for both H3K27ac and RNAP2-Ser5ph regions.
D. Distribution of average intensities in all signals for all H3K27ac and RNAP2-Ser5ph tracks (i.e. each track's signal is averaged, and contributes one data point to the distribution).

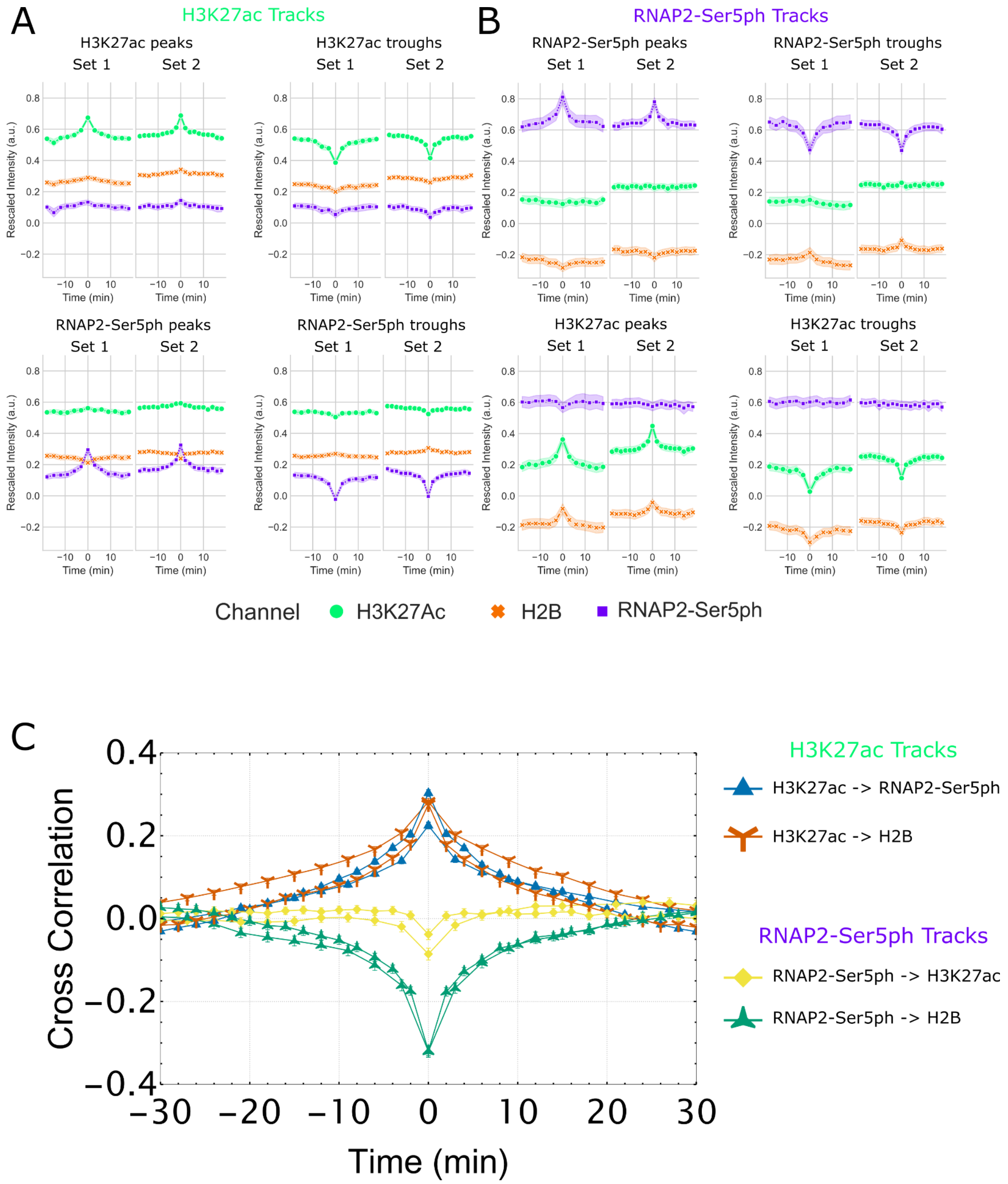


**Sup Fig. 4: Peak and Trough Detection Combined with Cross-Correlation Analysis Verify H3K27ac, RNAP2-Ser5ph and H2B Dynamic Behavior**

A. Detected peaks and troughs in all H3K27ac enriched tracks for two data sets of n=13 cells and n=15 cells. Both RNAP2-Ser5ph and H3K27ac peaks and troughs were detected, and all 3 signals were aligned based upon those peaks and troughs.
B. Same as (A), for RNAP2-Ser5ph enriched regions.
C. Cross correlation of various signals inside of H3K27ac and RNAP2-Ser5ph enriched regions, calculated for all tracks of length 20 frames or greater. Error bars are standard deviation obtained via bootstrapping (n=1000).

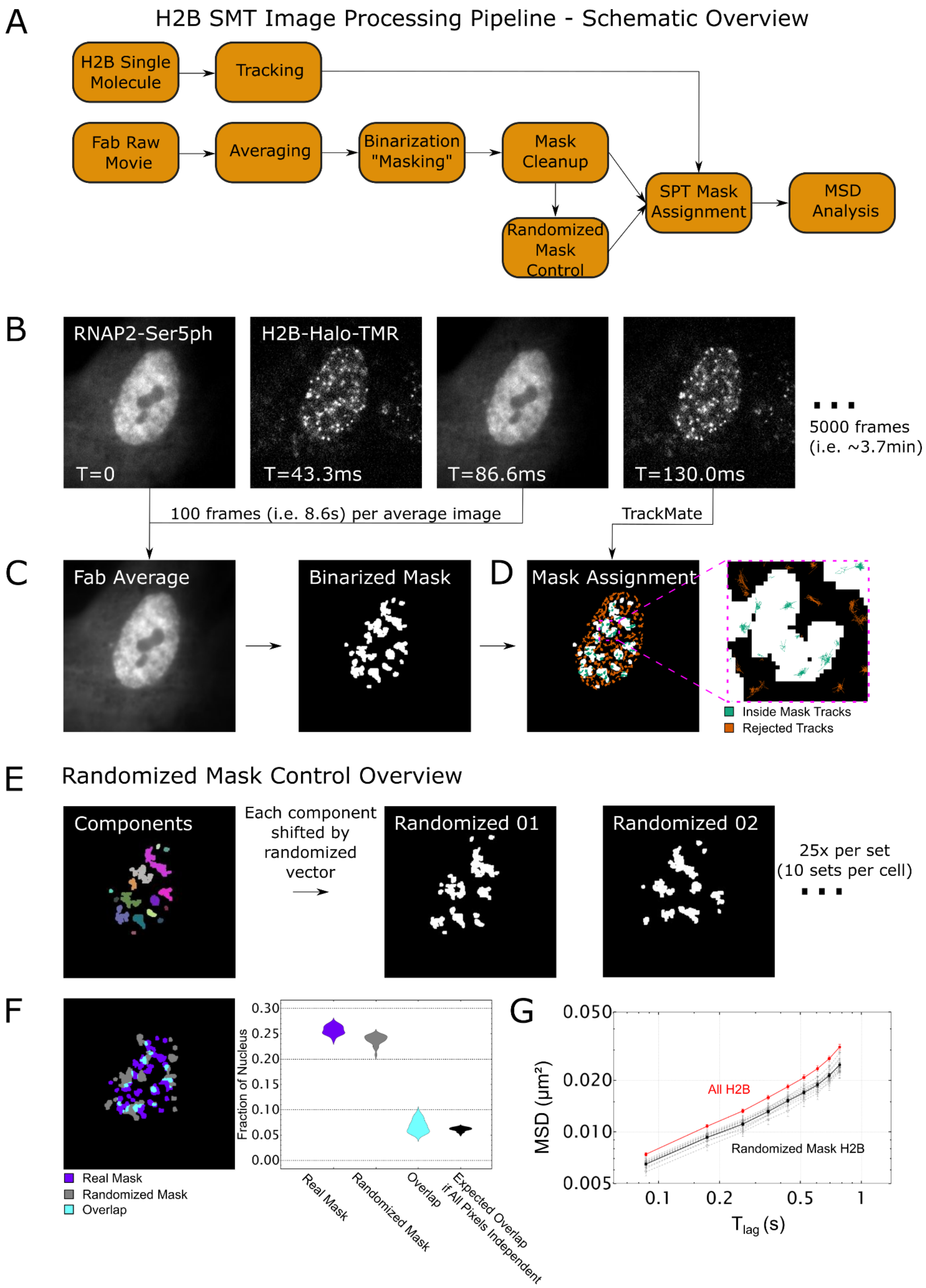
A
H2B SMT Image Processing Pipeline - Schematic Overview
H2B Single Molecule
Tracking
Fab Raw Movie
Averaging
Binarization "Masking"
Mask Cleanup
Randomized Mask Control
SPT Mask Assignment
MSD Analysis
B
RNAP2-Ser5ph
H2B-Halo-TMR
T=0
T=43.3ms
T=86.6ms
T=130.0ms
5000 frames (i.e. ~3.7min)
100 frames (i.e. 8.6s) per average image
TrackMate
C
Fab Average
Binarized Mask
D
Mask Assignment
Inside Mask Tracks
Rejected Tracks
E
Randomized Mask Control Overview
Components
Each component shifted by randomized vector
Randomized 01
Randomized 02
25x per set (10 sets per cell)
F
Real Mask
Randomized Mask
Overlap
Fraction of Nucleus
0.30
0.25
0.20
0.15
0.10
0.05
0.00
Real Mask
Randomized Mask
Overlap
Expected Overlap if All Pixels Independent
G
MSD (μm²)
0.050
0.020
0.010
0.005
All H2B
Randomized Mask H2B
0.1
0.2
0.5
1
T_lag (s)

**Sup Fig. 5: Analysis Pipeline for Tracking Single Nucleosomes in the Context of Post-Translational Modifications**

A. Schematic overview of the image processing pipeline for H2B single-molecule tracking experiments
B. Fab and single-molecule H2B are imaged in an alternating fashion, for 5000 total frames (3.7min).
C. 100 Fab images are used to create an average intensity image, reducing the noise generated by freely diffusing Fab. A local adaptive binarization algorithm is then applied to create a mask of enriched regions.
D. Determination is made whether single nucleosome tracks are inside or outside of the mask, and tracks outside of a mask are not used in subsequent analysis.
E. To create a randomized mask, first the connected pixels in a mask are segmented into separate regions, or “components”. Entire components are shifted by an individually randomized vector, creating a mask with random distribution of components, but whose morphology is analogous to the original mask.
F. Overlap between real and randomized masks. Distribution of overlap is quantified and compared to an *in-silico* control, the expected overlap between masks if all pixels in the nucleus were randomized individually. n=25 real and randomized masks
G. Log-log plot of all nuclear H2B MSD in a single cell vs those inside of 10 sets of 25 randomized masks for the same cell. The EA-TA-MSD for H2B inside randomized masks is always slightly decreased, demonstrating the need for a randomized mask control.
